# A Novel AI-Designed Antimicrobial Peptide Synergistically Potentiates Aminoglycosides against Colistin- and Carbapenem-Resistant *Acinetobacter baumannii*

**DOI:** 10.1101/2023.11.23.568446

**Authors:** Vipasha Thakur, Varsha Gupta, Prince Sharma, Anvita Gupta, Neena Capalash

## Abstract

The urgent necessity for new antibiotics becomes glaringly evident with the relentless rise of multidrug-resistant (MDR) *Acinetobacter baumannii* in clinical environments, where its infections lead to alarmingly high mortality rates. Antimicrobial peptides (AMPs) represent a promising novel option to combat nosocomial infections caused by MDR *A. baumannii*. In this study, six novel synthetic peptides were designed through generative artificial intelligence (AI) and synthesized for further experiments. Peptides AIG-R1, AIG-R4, and AIG-R5 showed potent broad-spectrum antibacterial activity against Gram positive and Gram negative pathogens. One of the peptides, AIG-R5, was effective even against colistin and carbapenem-resistant strains of *A. baumannii,* prevented biofilm formation, and eradicated established biofilms by 60%. Notably, AIG-R5 enhanced the activity of different antibiotics and was found to exhibit synergistic activity with antibiotics from the Aminoglycoside class. The combination of AIG-R5 and Tobramycin at 1/8×MIC and 1/4×MIC effectively reduced pre-formed biofilms of carbapenem resistant *A. baumannii* more than either component alone, as documented by confocal laser scanning microscopy (CLSM). Significant dose reduction and negligible cytotoxicity exhibited by AIG-R5 with aminoglycosides further encourages evaluation of the combination’s therapeutic potential *in vivo* against MDR *A. baumannii* infections.

## Introduction

*Acinetobacter baumannii* is an opportunistic nosocomial pathogen with the highest documented level of resistance in clinical settings, and many strains are incurable (Jung et al., 2021; Piperaki et al., 2019). The World Health Organization (WHO) and Centres for Disease Control and Prevention (CDC) have listed *A. baumannii* as a critical priority pathogen for which novel treatment options are urgently needed (WHO, 2017; CDC, 2019). *A. baumannii* (21.07%) and *Klebsiella pneumoniae* (29.03%) were among the most common bacterial pathogens isolated from COVID-19 patients (Vijay et al., 2021). Approximately 2% of all infections in health care settings are caused by *A. baumannii* in the United States and Europe, and this percentage is roughly twice as high in Asia and the Middle East (Colquhoun and Rather, 2020). Over the last decade, there has been a sharp increase in the prevalence of carbapenem-resistant *A. baumannii* (CRAB) (Su et al., 2022). Currently, colistin is the only antimicrobial peptide (AMP) approved by FDA that is active against MDR Gram negative bacteria (Costa et al., 2019). It is the last resort antibiotic used for the treatment of carbapenem-resistant *A. baumannii* infections as significant nephrotoxicity and neurotoxicity limit its use (Sacco et al., 2022). Moreover, colistin resistance has been reported in *A. baumannii* across all continents (Mirjalili et al., 2022; Upmanyu et al., 2022). Hence, there is a dire need of novel therapeutics to treat this extremely difficult drug-resistant pathogen that colonises and generates biofilms on biotic and abiotic surfaces, making eradication from clinical settings a significant challenge (Chapartegui-González et al., 2018).

Antimicrobial peptides (AMPs) are a promising alternative to traditional antibiotics due to their broad antibacterial activity and robustness against resistance development. AMPs composed of 11 to 50 amino acids are generally cationic peptides that interact with the anionic bacterial membrane via electrostatic interactions and lead to membrane disruption (Jaśkiewicz et al., 2019). A few AMPs have shown the ability to kill bacteria inside the biofilm and inhibit biofilm formation directly (Mazurkiewicz-Pisarek et al., 2023). However, these peptides have not been tested for synergy with existing antibiotics. When a single antibiotic fails and no new antibiotics are available, potentiation of already-existing drugs to treat *A. baumannii* infections assumes importance. AMPs are an effective substitute to conventional antibiotics when administered alone or in combination against multidrug resistant bacteria (Sacco et al., 2022). To effectively treat severe infections brought on by carbapenem and colistin resistant *A. baumannii*, a superior combination therapy must be developed.

Despite the advantages of AMPs, drawbacks such as high off-target toxicity, chemical and physical instability, and high production cost have been obstacles to the successful clinical development of AMPs (Chen and Lu, 2020). Given the increasing availability of data on AMP sequences, Artificial Intelligence (AI)-guided peptide design is an approach for creating new AMPs that are specifically designed to overcome these drawbacks.

Traditionally, Machine Learning (ML) models have been used to predict antimicrobial activity from libraries of existing molecules. Bidirectional long short-term memory (Bi-LSTM) recurrent neural networks have been applied to predict antimicrobial activity from a peptide’s sequence (Youmans et al., 2017). Other ML models have been used to predict peptide toxicity and stability (Khabbaz et al., 2021; Mathur et al., 2018).

On the other hand, Generative AI models can be used for *de novo* design of biological molecules, including small molecule drugs, DNA sequences, and new proteins (Gupta et al., 2018; Strokach and Kim, 2022; Taskiran et al., 2022; Wu et al., 2021). In particular, generative AI models can automate the process of designing new AMPs that do not exist in nature. Recurrent Neural Networks, Generative Adversarial Networks, and Variational Autoencoders have been used to *de novo* design AMP sequences of varying lengths (Dean and Walper, 2020 Gupta and Zou, 2019 Müller et al., 2018). However, wet lab validation has not been provided for many of these published models. In the few cases where the peptides have been tested experimentally, the AI-designed AMPs have not been evaluated or further optimized for important properties such as hemolytic activity, cytotoxicity, chemical and physical stability, and their ability to eradicate biofilms. Without studying these properties, these AI-designed AMPs face the same problems of toxicity and instability that plague natural AMPs. Using AI to design AMPs that can proceed to the clinic is complex and requires multiparameter optimization. Published models only consider how to design new peptide sequences with antibacterial activity.

Here, we present the first AI-designed peptide, AIG-R5, with broad spectrum antibacterial activity against MDR pathogens, low off-target hemolysis, the ability to inhibit and eradicate biofilms of MDR *A. baumannii*, and synergy with an existing class of antibiotics: the aminoglycosides. In this study, six novel peptides AIG-R1 to AIG-R6 were designed using generative AI models, and their therapeutic properties were predicted using a suite of ML models. These peptides were chemically synthesized and evaluated for antimicrobial activity against Gram positive and Gram negative pathogens of ESKAPE group, including colistin (CoR) and carbapenem resistant *A. baumannii* (CRAB) strains. Of these six peptides, the most effective peptide AIG-R5 was found to have broad spectrum of activity and was effective against CoR and CRAB strains of *A. baumannii.* This peptide significantly inhibited biofilm formation of MDR *A. baumannii*, eradicated preformed biofilms, and potentiated the activity of conventional antibiotics against *A. baumannii*.

## Results

### 1.1 Peptide Design and Synthesis

AIG-R1 was the first peptide designed using AIGenus^®^, the generative artificial intelligence platform for protein design (AINovo Biotech). AIG-R2 through R6 are derivatives of AIG-R1 which were designed through our AI-guided iterative mutational approach to reduce off-target toxicity and increase stability. All peptides’ antimicrobial activity, hemolytic activity, and stability were predicted through several machine learning models in the AIOptimus^®^ suite, and physiochemical properties such as hydrophobic moment and charge were also calculated from the amino acid sequence (Table 1). While AIG-R1 had predicted probability of 0.604 of being non-hemolytic, the top performing derivative peptides, AIG-R4 and AIG-R5, had increased non-hemolytic probabilities of 0.794 and 0.840, respectively. In addition, the predicted probability of the peptide being stable (having a half-life greater than 45 minutes) increased from 0.846 in AIG-R1 to above 0.94 for AIG-R2 through AIG-R6. At the same time, the predicted probability that the peptide is active against *A. baumannii* (MIC <128µM) remained above 0.99 for AIG-R2 through AIG-R6. The peptides were synthesized chemically with >90% purity as measured by HPLC. This AI-based approach allowed for the rapid generation and screening of millions of peptide sequences, enabling the identification of new peptides with high bactericidal activity against target pathogens.

**Table 1.**
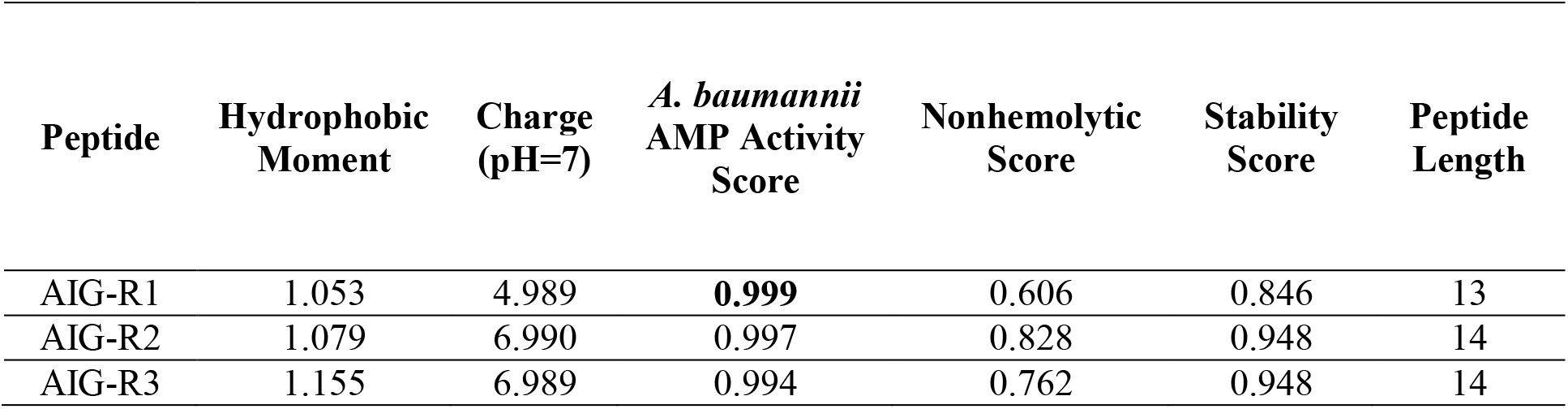
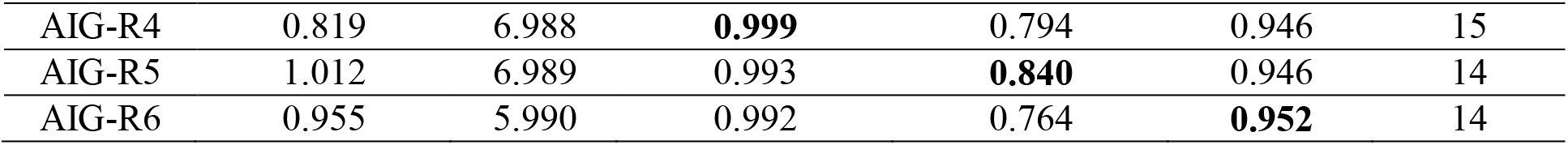
Physiochemical properties and predicted therapeutic properties of the six AI-designed peptides.

### 1.2 Antimicrobial activity

Of the six peptides tested, AIG-R5 exhibited the strongest antibacterial activity against *A. baumannii* ATCC 17978 with MIC of 2µM. For AIG-R1 and AIG-R4, the MIC was 4 and 3µM, respectively. MIC of 20 µM was observed with AIG-R2 and AIG-R3; for AIG-R6 the MIC was 22µM. For all peptides, the experimental MIC against *A. baumannii* was less than 128µM, validating the predictions by the AIOptimus^®^machine learning models. AIG-R1, AIG-R4 and AIG-R5 were found to be the most effective against *A. baumannii* and other ESKAPE group pathogens (Table 2). Therefore, AIG-R1, AIG-R4 and AIG-R5 were selected for further studies.

**Table 2.** MIC of peptides against Gram positive and Gram negative bacteria of the ESKAPE group of pathogens.

| Microorganism | Strain | AIG-R1 | AIG-R2 | AIG-R3 | AIG-R4 | AIG-R5 | AIG-R6 |
| --- | --- | --- | --- | --- | --- | --- | --- |
| *Acinetobacter baumannii* | ATCC 17978 | 4 | 20 | 20 | 3 | 2 | 22 |
| *Pseudomonas aeruginosa* | PAO1 | 6 | 8 | >18 | 6 | 6 | >18 |
| *Klebsiella pneumonia* | KP-1 | 4 | 14 | >14 | 3 | 4 | >16 |
| *Staphylococcus aureus* | ATCC 25923 | 4 | 12 | 10 | 3 | 4 | 12 |
| *Enterococcus* spp. | B-3650 | 12 | >8 | >12 | 10 | >8 | >12 |

AIG-R1, AIG-R4 and AIG-R5 were equally effective against colistin and carbapenem resistant *A. baumannii* strains (Table 3). Antibiotic resistance profiles of these isolates are given in supplementary table S2.

**Table 3.** MIC of antimicrobial peptides against colistin and carbapenem resistant *A. baumannii* (CRAB) strains.

| S. No. | *A. baumannii* | MIC (µM) of AIG- | MIC (µM) of | MIC (µM) of |
| --- | --- | --- | --- | --- |

|  | strains | R1 | AIG-R4 | AIG-R5 |
| --- | --- | --- | --- | --- |
| 1. | Ci3 | 4 | 3 | 2 |
| 2. | Ci6 | 4 | 3 | 2 |
| 3. | Ci7 | 4 | 3 | 2 |
| 4. | P-1139 | 4 | 3 | 4 |
| 5. | P-1182 | 4 | 3 | 4 |
| 6. | P-1119 | 4 | 3 | 2 |
| 7. | P-1131 | 4 | 3 | 2 |
| 8. | P-1169 | 4 | 3 | 2 |
| 9. | DHA 8 | 4 | 3 | 2 |
| 10. | DHA 10 | 4 | 3 | 2 |

### 1.3 Stability of antimicrobial peptides

No loss in activity of AIG-R5 was observed from 3 to 12 pH as the MIC remained unchanged against *A. baumannii* 17978. However, the activity of AIG-R1 and AIG-R4 decreased with an increase in pH. At pH of 12, there was a 2-fold and 3-fold reduction in activity of AIG-R4 and peptide AIG-R1, respectively (Supplementary table S3).

Further, the stability of AMPs was investigated at the physiological concentration of Na^+^, Mg^2+^ and Ca^2+^ ions. The activity of AIG-R1 remained unchanged in presence of these cations. The antimicrobial activity of AIG-R4 also remained unchanged in presence of Na^+^ ions, but the MIC increased by 1.6 and 2-fold in the presence of Mg^2+^ and Ca^2+^ respectively. There was a change in the MIC of AIG-R5 from 2µM to 4µM in the presence of physiological concentration of Na^+^ and Mg^2+^ ions. However, no loss in activity was observed in the presence of Ca^2+^ ions. There was reduction in the activity of AIG-R1 (2-3 fold), AIG-R4 (2-4 fold) and AIG-R5 (4-6 fold) in the presence of 3 and 5% serum (Table 4).

**Table 4.** Stability of AMPs AIG-R1, AIG-R4, and AIG-R5.

| Peptide | MIC (µM) against *A. Baumannii* 17978 | MIC (µM) in presence of 150mM NaCl | MIC (µM) in presence 1mM MgCl<sub>2</sub> | MIC (µM) in presence 2.5mM CaCl<sub>2</sub> | MIC (µM) in presence of 3% serum | MIC (µM) in presence of 5% serum |
| --- | --- | --- | --- | --- | --- | --- |
| AIG-R1 | 4 | 4 | 4 | 4 | 8 | 12 |
| AIG-R4 | 3 | 3 | 5 | 6 | 6 | 12 |
| AIG-R5 | 2 | 4 | 4 | 4 | 8 | 12 |

### 1.4 Hemolytic and Cytotoxic activity

The hemolysis of human RBCs was assessed at different concentrations of AMPs. A concentration dependent increase in hemolysis was observed. Hemolysis was <5% at the MIC of AIG-R1, AIG-R4 and AIG-R5. However, at 16×MIC the hemolysis increased to 88% with AIG-R1. The hemolysis at 16xMIC was reduced in AIG-R5 and AIG-R4, with 38% hemolysis observed with 16xMIC of AIG-R5 and 10% with 16xMIC of AIG-R4 (Figure 1a).

**Figure 1.**
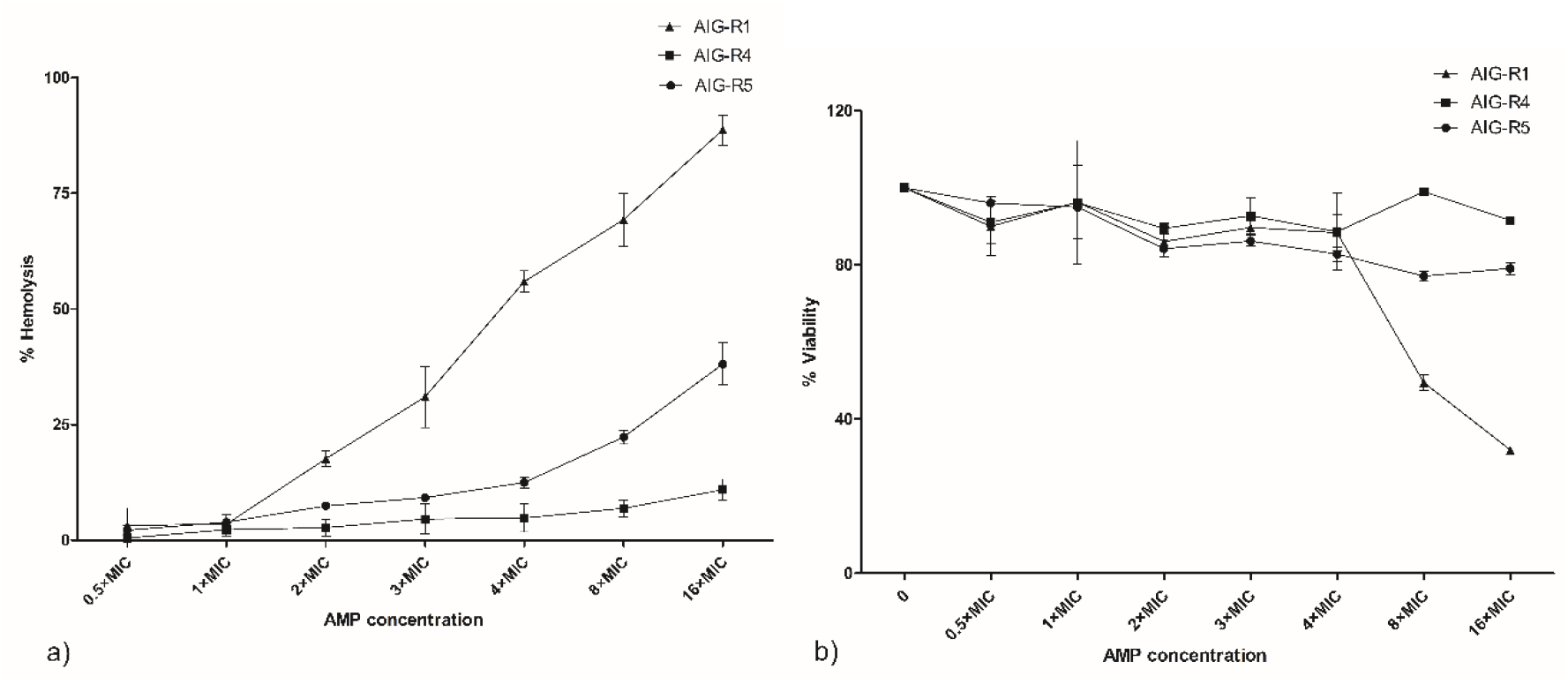
**(a)** Hemolytic activity of AMPs against human RBCs. (b) Cytotoxicity of AMPs against HeLa cell line. All values expressed as the mean ± SD of three independent experiments.

The cytotoxicity of the three peptides was investigated against the HeLa cell line. More than 95% cell survival was observed at MIC with all three peptides. Even though AIG-R1 showed potent antibacterial activity, this peptide demonstrated high hemolysis and cytotoxicity at higher concentrations. Derivatives AIG-R4 and AIG-R5 were less cytotoxic, with cell viability of 91.5 and 79% at 16×MIC, respectively, in comparison to AIG-R1 where cell viability fell to 32% at 16xMIC (Figure 1b).

### 1.5 Membranolytic activity

*A. baumannii* cells showed outer membrane damage as reflected by NPN uptake upon treatment with peptides. The maximum uptake of NPN was observed with AIG-R5, followed by AIG-R1 and AIG-R4 after 30 minutes of treatment (Figure 2a). Treatment at 1× and 2×MIC for 5 minutes also showed inner membrane depolarisation as revealed by DiOC_2_ (3,3’-Dimethyloxacarbocyanine), a membrane potential-sensitive dye (Figure 2b).

**Figure 2.**
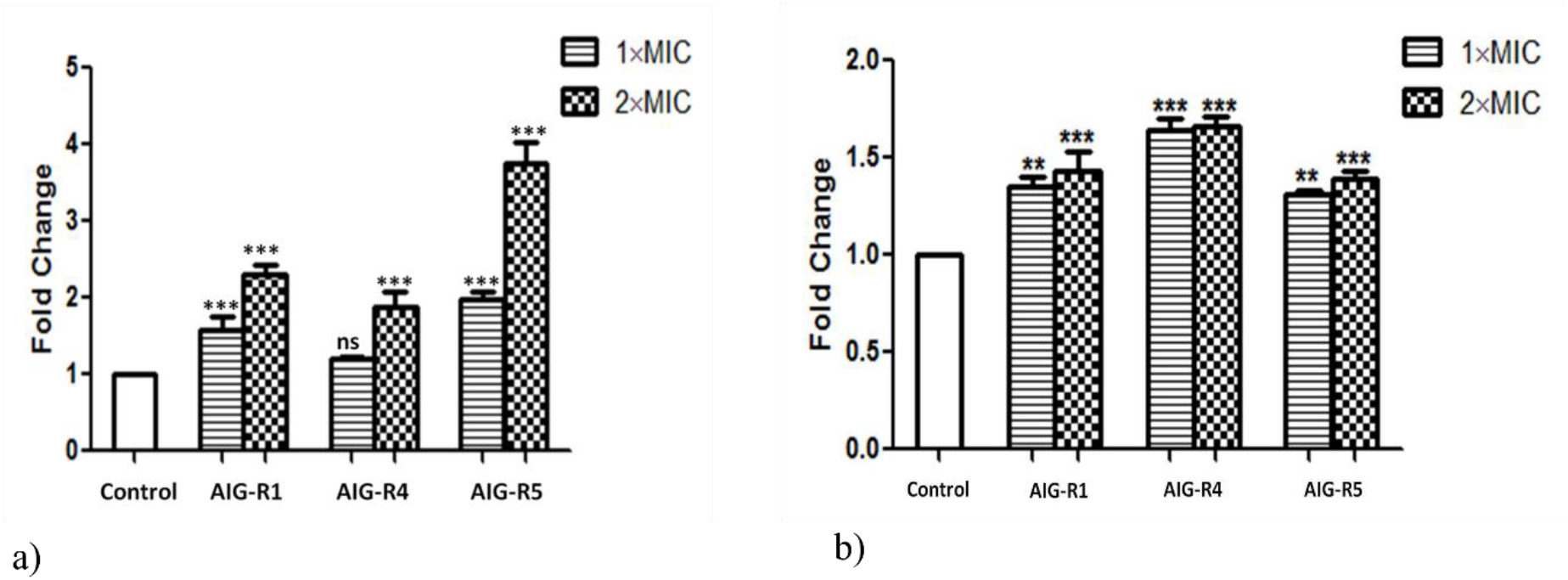
(a) Outer membrane permeability as determined by NPN uptake, and (b) Inner membrane depolarization of *A. baumannii* ATCC 17978 as measured by the membrane potential-sensitive dye, DiOC_2_ after treatment with peptides for 5 minutes. All results presented as mean ± SD of three independent experiments. \*\*\**p*<0.001, \*\**p<*0.01, ns - not significant compared to control.

### 1.6 Reactive Oxygen Species (ROS) production

*A. baumannii* cells showed maximum ROS production after treatment with AIG-R1 (3.6-fold) at 1×MIC followed by AIG-R4 (2.3-fold) and least (1.2-fold) with AIG-R5 (Figure 3). However, the pre-treatment with sub-inhibitory concentration of ROS quenchers, thiourea and 2.2’-bipyridyl (Supplementary Table S1) did not affect the antimicrobial activity of the peptides. This suggested that ROS generation was not responsible for their antimicrobial activity.

**Figure 3.**
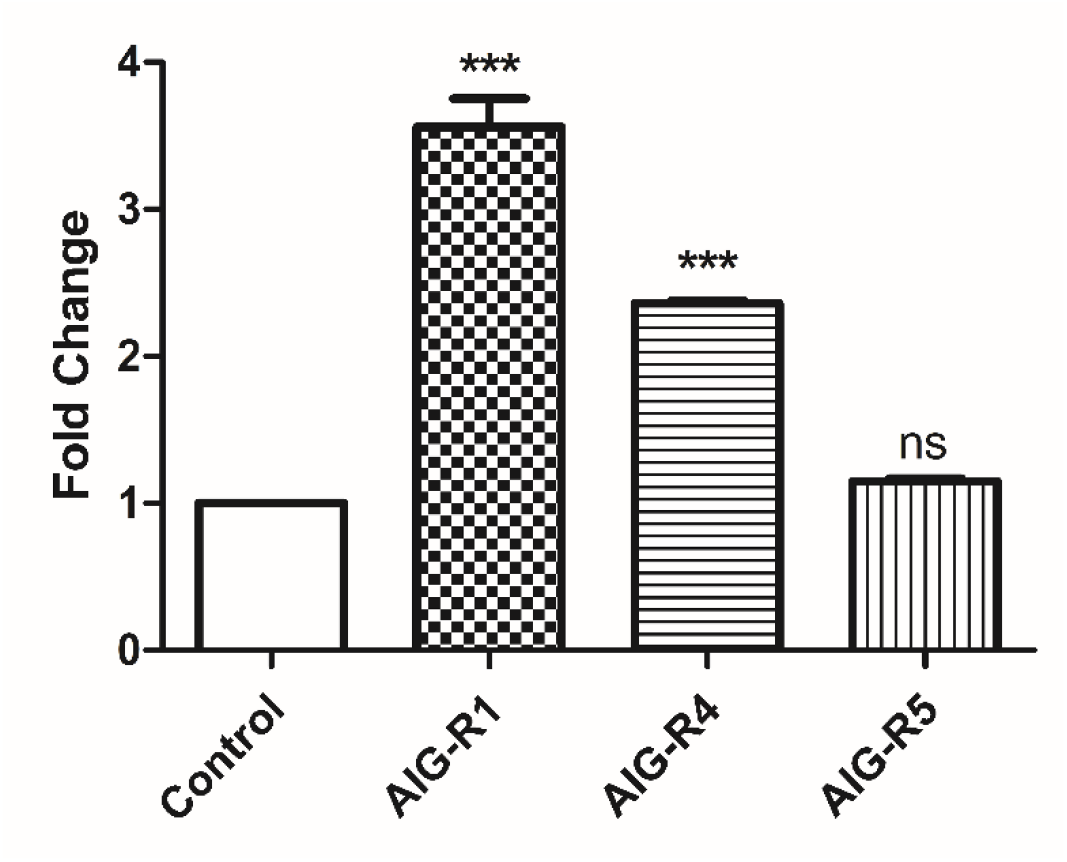
ROS production in *A. baumannii* ATCC 17978 cells on treatment with AIG-R1, AIG-R4 and AIG-R5 at 1×MIC for 60 minutes. \*\*\**p*<0.001, ns - not significant, compared to control. Data presented as mean ± SD of three independent experiments.

### 1.7 Combination of peptides with antibiotics

AIG-R1 was found to exhibit an additive effect (FIC 0.5 to 1) with different classes of conventional antibiotics against *A. baumannii*. AIG-R4 demonstrated synergistic activity (FIC≤0.5) when combined with colistin, and showed an additive effect with imipenem, rifamycin, ciprofloxacin, tobramycin, amikacin and kanamycin. AIG-R5 demonstrated synergistic activity with antibiotics from the aminoglycoside class, including Tobramycin, Amikacin, and Kanamycin Sulphate (Table 5). The combination of 1/8×MIC (0.25µM) of AIG-R5 with 1/4×MIC (2µg/ml) Tobramycin was synergistically effective (FIC<0.5) against *A. baumannii* 17978 which was corroborated by FESEM analysis after treatment. The treatment of cells with this combination resulted in the loss of cellular integrity and membrane rupture (Figure 4). This combination was also found to be effective against the colistin and carbapenem resistant *A. baumannii* strain DHA-10.

**Figure 4.**
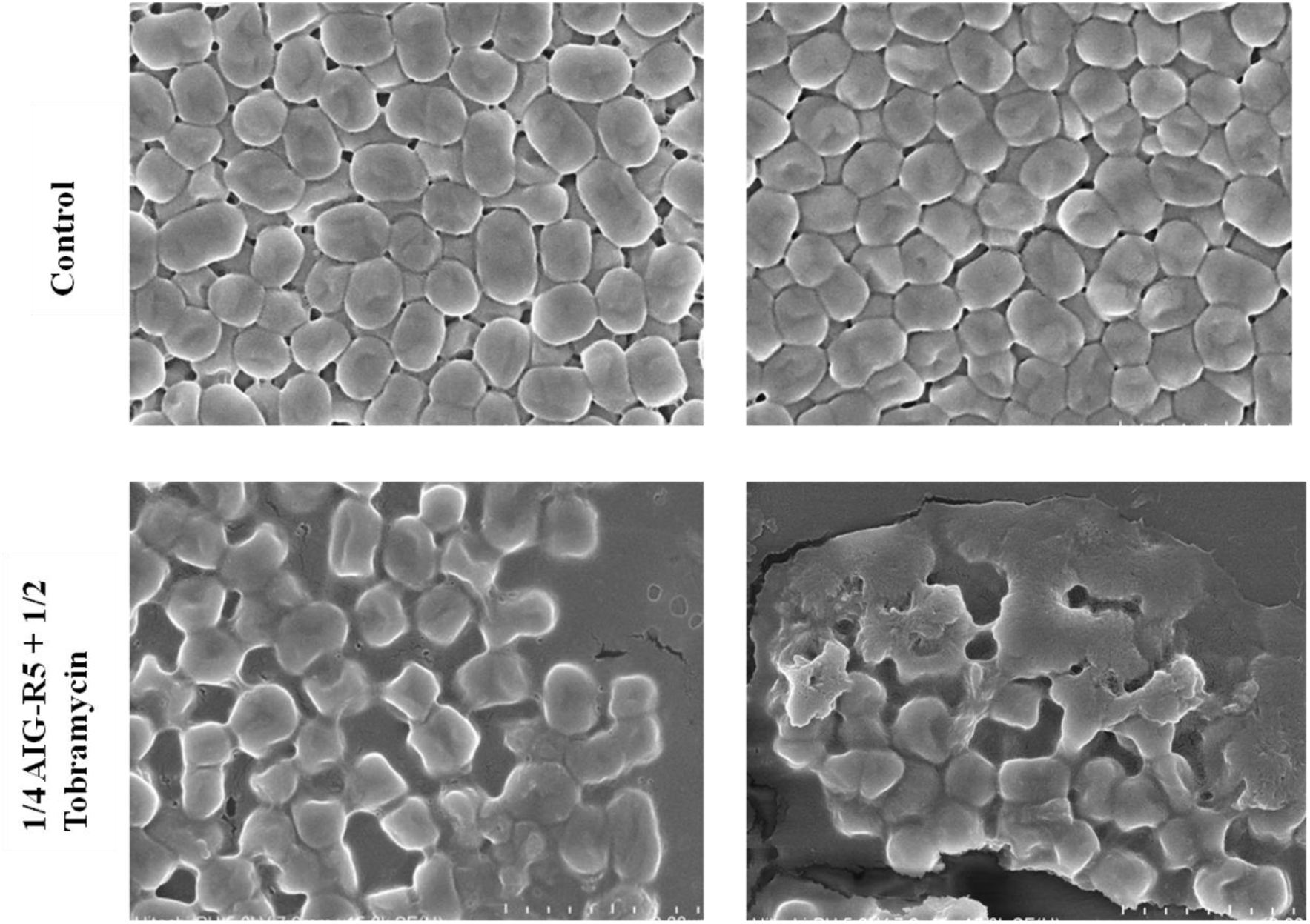
FESEM analysis of 2×FIC (1/4×MIC AIG-R5 AND 1/2×MIC tobramycin) treated and untreated *A. baumannii* ATCC 17978 cells.

**Table 5.**
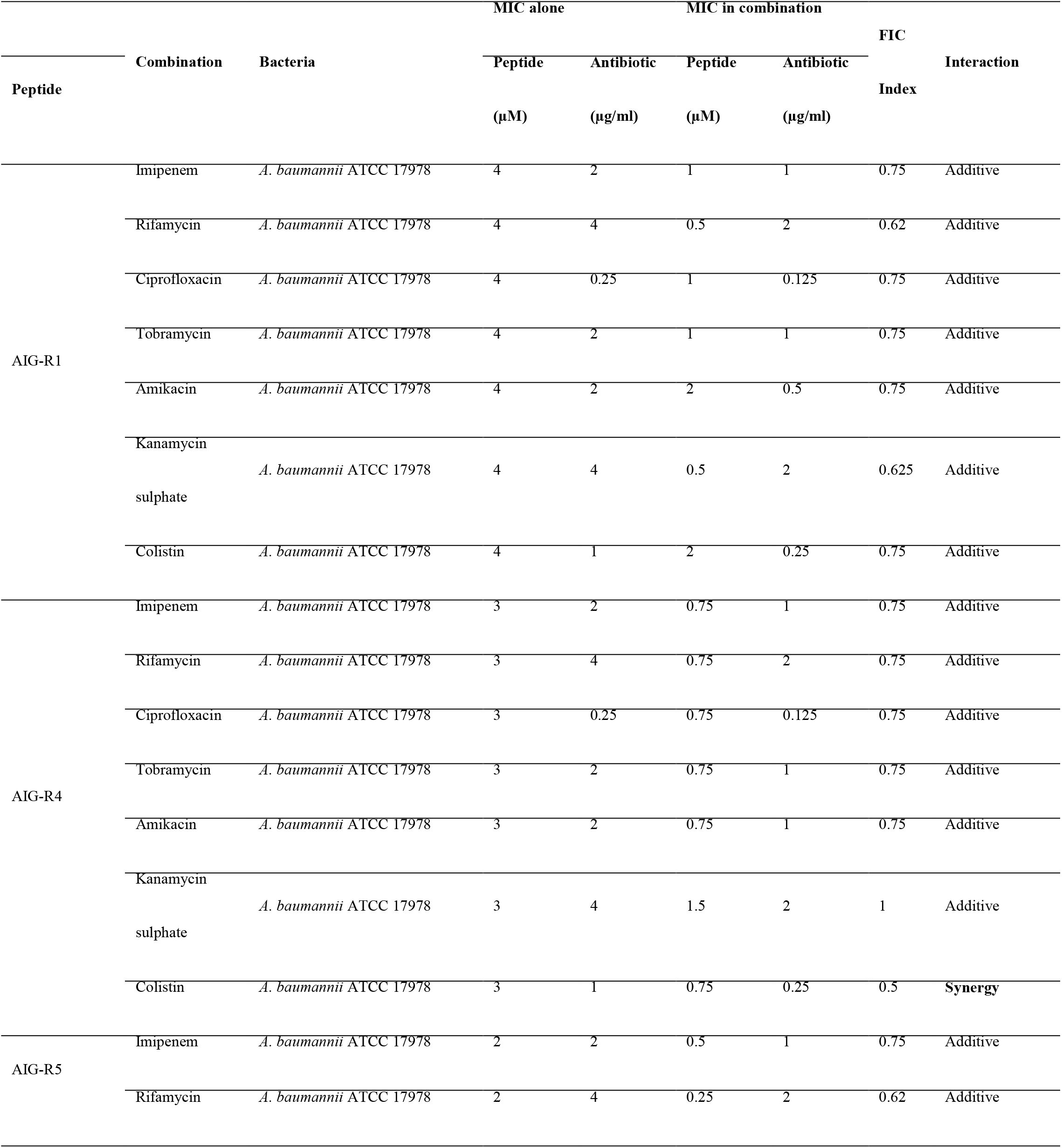
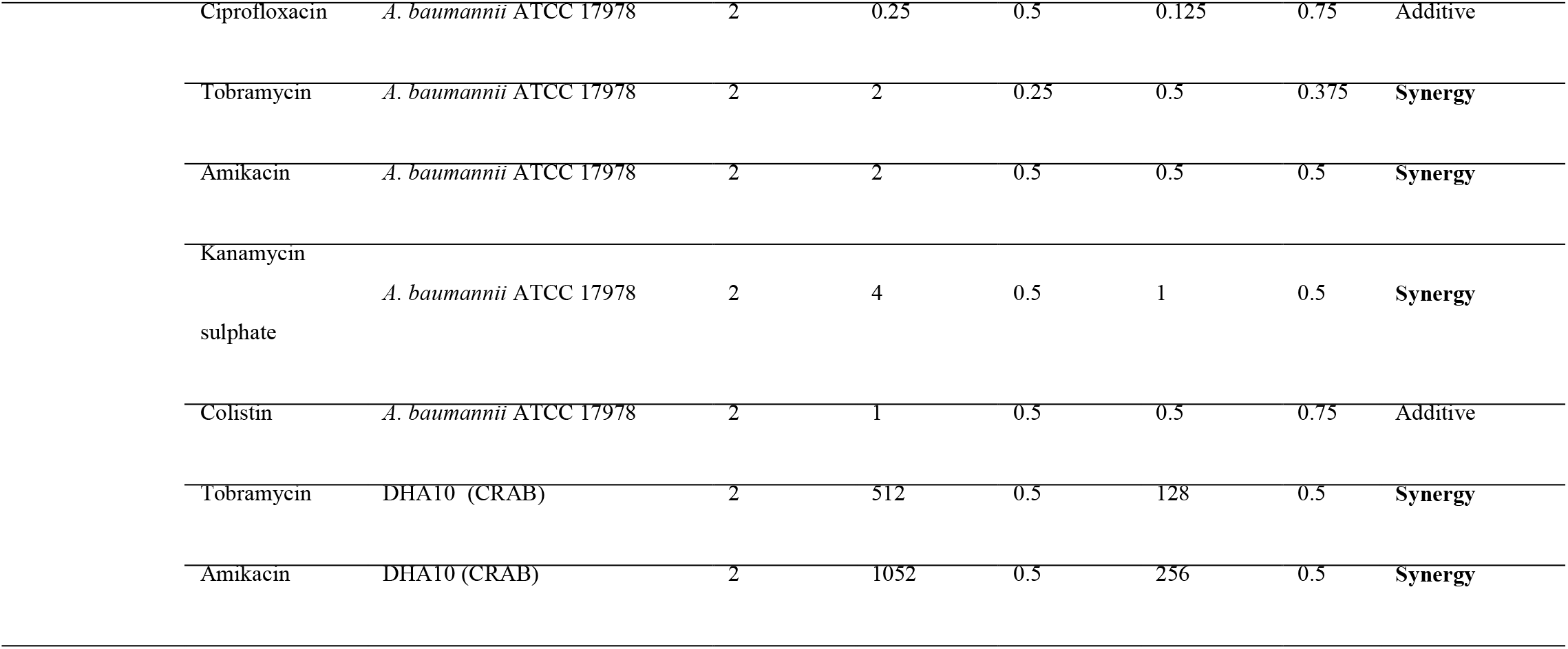
Interaction of AMPs with different antibiotics.

### 1.8 Biofilm Inhibition

Since AIG-R5 significantly potentiated the activity of antibiotics against *A. baumannii,* it was further evaluated for its effect on biofilm formation and eradication, alone and in combination with antibiotics. AIG-R5 inhibited biofilm formation in *A. baumannii* 17978 in a concentration dependent manner with 25, 32 and 60% reduction at 2 (1×MIC), 4 (2×MIC) and 8µM (4×MIC), respectively in comparison to the control. Similarly, there was decrease in biofilm formation in colistin and CRAB strains (P-1139 and DHA10), which otherwise formed a good biofilm (Supplementary Figure S1). The reduction of 25, 33 and 58% was observed in biofilm of DHA10 at 2 (1×MIC), 4 (2×MIC) and 8µM (4×MIC) concentrations of AIG-R5. Similarly, biofilm inhibition of 23, 35 and 54 % was observed in P-1139 (Figure 5a).

**Figure 5.**
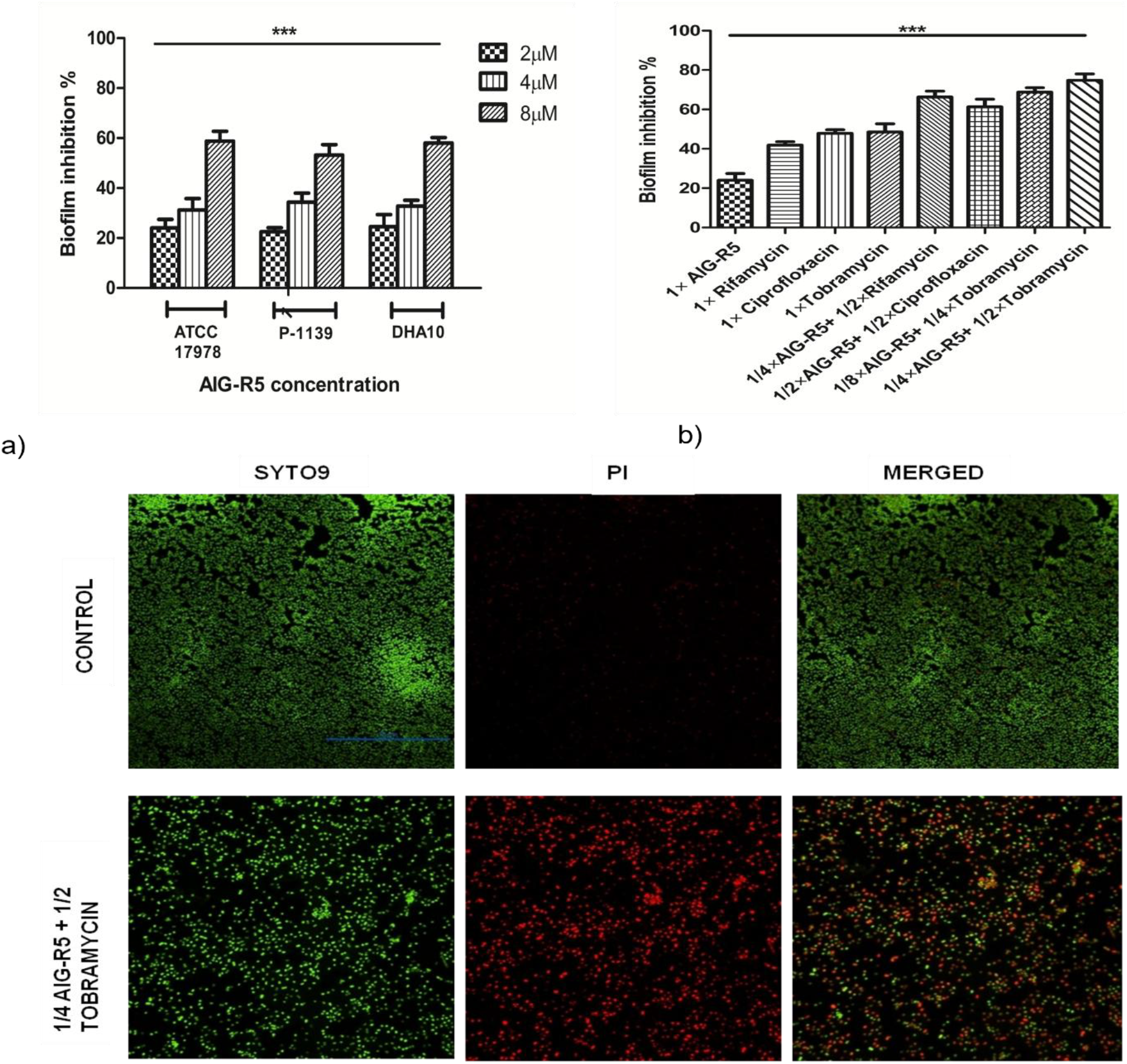
**(a)** Biofilm inhibition in *A. baumannii* ATCC 17978 and colistin and CRAB strains (P-1139 and DHA10) in the presence of AIG-R5, **(b)** Biofilm inhibition in *A. baumannii* ATCC 17978 in presence of AIG-R5, antibiotics alone (rifamycin, ciprofloxacin and tobramycin) and combination of AIG-R5 with antibiotics. **(c)** CLSM images showing *A. baumannii* 17978 biofilm inhibition in presence of 1/4×MIC (0.5µM) AIG-R5 and 1/2×MIC (1µg/ml) tobramycin after staining with SYTO9 and Propidium iodide. All values expressed as the mean ± SD of three independent experiments.

The combination of AIG-R5 and tobramycin at 1× and 2×FIC inhibited biofilm formation in *A. baumannii* 17978 by 70 and 75%, respectively, which was significantly higher than AIG-R5 or tobramycin used alone (Figure 5b). Co-treatment with ¼×MIC (0.5µM) of AIG-R5 with ½×MIC (2µg/ml) rifamycin and ¼×MIC (0.5µM) of AIG-R5 with ½×MIC (0.125µg/ml) ciprofloxacin also led to 67% and 62% inhibition of biofilm formation.

CLSM confirmed antibiofilm activity on live/dead staining with SYTO9 and Propidium iodide (PI). One day old, untreated *A. baumannii* biofilm was dense showing live bacteria emitting green fluorescence. The biofilm inhibition observed at 1×FIC (1/8×AIG-R5+ 1/4×Tobramycin) showed dispersed cells (Supplementary Figure S2). However, the biofilm formed in presence of 2×FIC (1/4×AIG-R5+ 1/2×Tobramycin) revealed much scattered cells that were dead, exhibiting red fluorescence (Figure 5c). Quantitative analysis of CLSM images revealed that treated biofilm had 2.6-fold reduced proportion of live to dead cells.

### 1.9 Biofilm eradication

AIG-R5 eradicated 24-hour pre-formed biofilms of *A. baumannii* ATCC 17978, as well as colistin and CRAB strains (P-1139 and DHA10). As shown in Figure 5a, AIG-R5 at 16µM (8×MIC) eradicated up to 58% biofilm formed by *A. baumannii* 17978 and DHA10, while the peptide eradicated 45% in case of P-1139. However, the combination of AIG-R5 and tobramycin at 1×FIC and 2×FIC, even at a very low concentration, was able to eradicate the pre-formed biofilm of *A. baumannii* 17978 by 40 and 55%, respectively (Figure 6b). Moreover, the combination at 1×FIC and 2×FIC showed negligible cytotoxicity (Supplementary Figure S3, Table S4)

**Figure 6.**
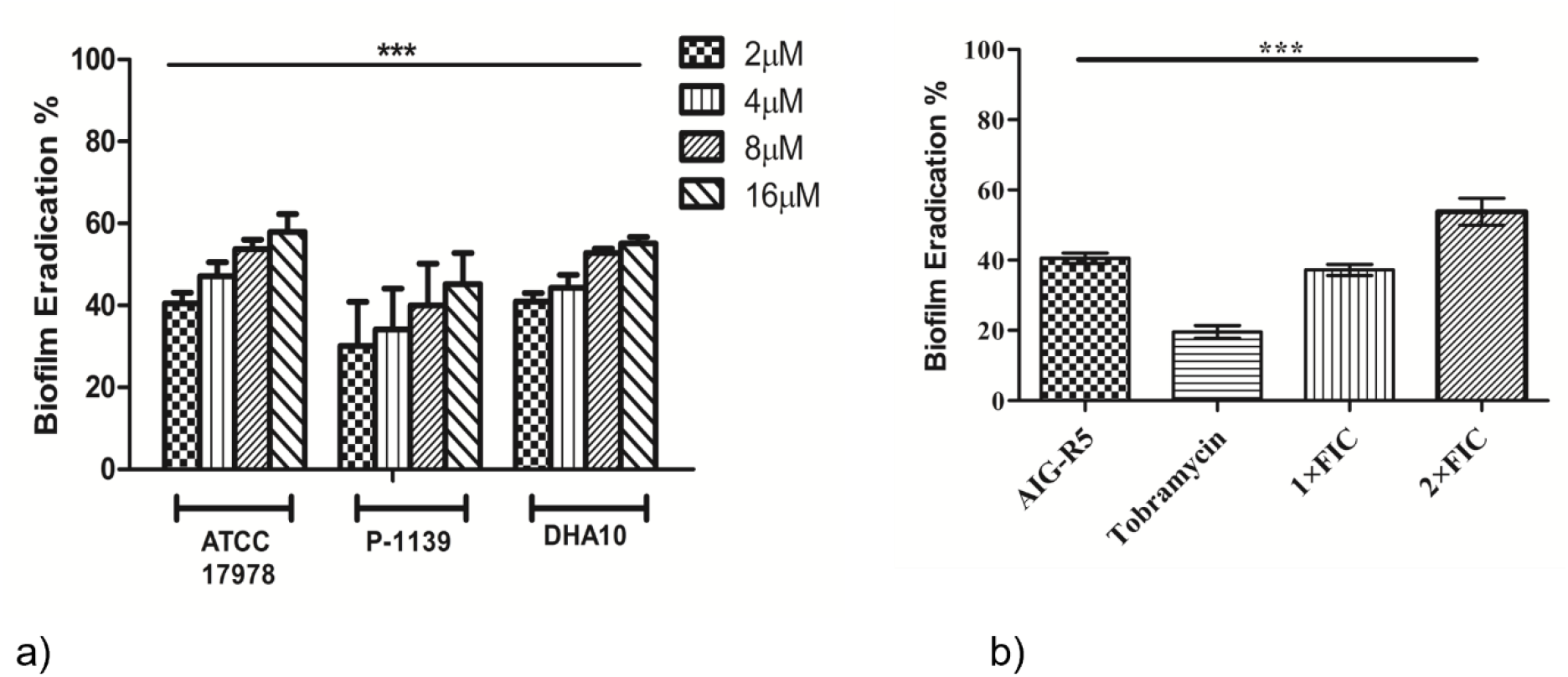
**(a).** Eradication of pre-formed biofilms of *A. baumannii* ATCC 17978 and colistin and CRAB strains (P-1139 and DHA10). (b) The eradication of pre-formed biofilm of *A. baumannii* ATCC 17978 by combination of AIG-R5 and tobramycin at 1× and 2×FIC. All values expressed as the mean ± SD of three independent experiments.

## Discussion

Bacterial infections caused by MDR bacteria are one of the most serious problems faced by clinicians worldwide (Choi et al., 2021). In this regard, *A. baumannii* is one of the most challenging pathogens among the ESKAPE group of drug-resistant pathogens, and the rise in carbapenem resistance is a great concern for public health (Nguyen and Joshi, 2021; Xie et al., 2018). Therefore, development of novel therapeutics against carbapenem resistant *A. baumannii* is a priority as per the orld Health Organization’s guidelines.

The present study reports the antibacterial activity of novel AI-designed synthetic peptides with a broad range of activity against Gram positive and Gram-negative pathogens of the ESKAPE group. AMPs AIG-R1, AIG-R4 and AIG-R5 were found to have outstanding activity against colistin and carbapenem resistant *A. baumannii,* and AIG-R5 was the most effective with an MIC of 2µM. These peptides acted by increasing the outer membrane permeability and disrupting the inner cell membrane of the bacteria. While the first synthesized peptide AIG-R1 also displayed an excellent MIC of 4 µM, this peptide had high cytotoxicity and hemolysis on human RBCs. Through AI-guided optimization, derivatives AIG-R4 and AIG-R5 were developed. Corresponding with *in silico* predictions, both peptides showed lower hemolysis and cytotoxicity than AIG-R1 while maintaining excellent broad-spectrum antibacterial activity.

The therapeutic potential of these peptides was evaluated in combination with conventional antibiotics used against *A. baumannii.* AIG-R5 synergistically potentiated the activity of aminoglycosides against *A. baumannii*. Through their distinctive membrane-disruption mechanisms, AMPs raise the permeability of anionic cell membranes, improve the entry of antibiotics and raise the sensitivity of cells to antibiotics (Lin et al., 2021). *A. baumannii* strains resistant to aminoglycosides such as amikacin, tobramycin and isepamicin have been reported worldwide (Upadhyay et al., 2018). The mechanism of resistance to aminoglycosides commonly involves the production of aminoglycoside modifying enzymes (AMEs) like aminoglycoside phosphotransferase (APT), aminoglycoside acetyltransferase (AAC) and aminoglycoside nucleotidyl transferase (ANT) (Laws et al., 2019). Cationic AMPs indolicidin and its analogs have been reported to inhibit aminoglycoside acetyltransferase and aminoglycoside phosphotransferase by binding to both the free enzyme and enzyme-substrate complexes (Boehr et al., 2003). Therefore, we hypothesize that the synergy of AIG-R5 with aminoglycoside class of antibiotics may be mediated through the inhibition of AMEs.

Biofilm development is a significant contributor to persistent and recurring infections (Dostert et al., 2019). Biofilm-related diseases, such as those spread by medical devices, increase the risk of infection and patient mortality (Mwangi et al., 2019). Biofilms typically require tenfold to 1000-fold higher concentration of conventional antibiotics for eradication due to the extracellular lipopolysaccharide matrix that obstructs antibiotic penetration (Ajish et al., 2022). AMPs exhibit anti-biofilm activity by forming pores within the lipid components of the biofilm, passing through the extracellular biofilms, or dispersing the biofilms. Although the exact mechanism of antibiofilm activity is not well established, the synergistic action of AMP and antibiotics may be efficient in breaking the biofilm matrix and allowing access to the bacterial cells within the biofilm (Chung and Khanum, 2017). AIG-R5 inhibited biofilm formation in *A. baumannii* in a dose dependent manner. It was able to suppress the *A. baumannii* biofilm formation at concentration lower than its MIC when combined with commonly used antibiotics including rifamycin, ciprofloxacin and tobramycin. The combination of AIG-R5 and tobramycin was effective even at concentration of 0.25µM (1/8×MIC) and 0.5mg/ml (1/4×MIC) in inhibiting biofilm formation. CLSM also corroborated a significant increase in the proportion of dead cells in biofilm treated with tobramycin and AMP as compared to the untreated biofilm. In addition, the combination of the peptide and antibiotic displayed negligible cytotoxicity, supporting its suitability for *in vivo* use.

The majority of AMPs display stronger anti-biofilm activity than biofilm eradication activity (Verderosa et al., 2019). Natural AMP LL-37 inhibited biofilm formation and eradicated pre-formed *A. baumannii* biofilms, but at very high concentration of 512µg/ml, suggesting that it would be more suitable for prevention of biofilm-associated infections than for their treatment (Jaśkiewicz et al., 2019). Marsupial AMP WAM-1 was effective at dispersing biofilms at 111µg/ml when used against clinical isolates, and therefore therapeutically was more potent than LL-37 (Spencer et al., 2018). Another AMP, magainin 2, degraded only 33.3% of pre-formed biofilms at 128µM (64×MIC) (Kim et al., 2018). However, the peptide AIG-R5 eradicated pre-formed *A. baumannii* ATCC 17978 biofilms by more than 50% at just 16µM (8×MIC) and was equally effective on pre-formed biofilms of colistin-resistant and CRAB isolates. The combination of tobramycin (0.5mg/ml, 1/4×MIC) and AIG-R5 (0.25µM, 1/8×MIC) even at a very low concentration was able to eradicate a pre-formed biofilm by 40%.

By using a lower concentration of the peptide with approved antibiotics in such a combination therapy, the toxicity and cost of treatment can be reduced, which are major deterrents that at present discourage the use of peptides for bacterial infections. To our knowledge, AIG-R5 is the first AI-designed peptide that shows broad-spectrum antibacterial activity and the ability to synergistically potentiate an existing class of antibiotics against MDR *A. baumannii*. AIG-R5 overcomes the issues of cost, toxicity, and stability that AMPs have faced in the past. Hence, the combination therapy of AIG-R5 and conventional antibiotics, especially the aminoglycosides, is being evaluated *in vivo* against carbapenem resistant *A. baumannii* in a pneumonia mouse infection model.

## Materials and Methods

### 1.10 Design and Synthesis of Peptides through Generative AI Models

To generate novel AMP sequences, we applied AIGenus^®^, a deep generative neural network for protein design developed by AINovo Biotech, Inc (“AINovo”). Briefly, a sequence-based generative model was pretrained on 100,000 helical peptides. The model was then fine-tuned on published data of experimentally validated AMPs (Jhong et al., 2019; Lee et al., 2015; Novković et al., 2012; Wang et al., 2016). After training, a library of 2000 novel AMP sequences was sampled from the trained generative model. AIOptimus^®^, a suite of machine learning models for virtual screening from AINovo, was used to predict antimicrobial activity against *A. baumannii* for all peptides in the generated library. Physiochemical properties such as hydrophobic moment and charge were calculated for each peptide using the modlAMP package in Python. Peptides were clustered based on their sequences and ranked by their predicted antimicrobial activity. The top peptide, AIG-R1, was synthesized and tested experimentally for antibacterial activity, hemolysis against human red blood cells (RBCs), and stability in human serum.

To reduce toxicity and improve stability, variants of AIG-R1 were designed by AINovo using a proprietary iterative mutation approach. In short, a large language model was trained to predict likely AMP mutations. Guided by this model, libraries of mutants of AIG-R1 were generated and evaluated *in silico* using machine learning classifiers from the AIOptimus^®^ suite. Sequences were filtered and ranked based on predicted antimicrobial activity, off-target toxicity, and stability. Active AMPs were defined as having MIC less than a cut-off of 128µM against *A. baumannii*. At the end of the process, the top five peptide sequences, AIG-R2 through R6, were chosen for synthesis.

### 1.11 Antimicrobial activity

The Minimum Inhibitory Concentration (MIC) of the six novel synthetic peptides was assessed against ESKAPE group pathogens using the broth microdilution method in Mueller Hinton (MH) broth (Weinstein, 2018). As per CLSI guidelines (2018), overnight grown culture was diluted in Mueller Hinton (MH) broth to a turbidity equivalent to 0.5 McFarland standards. Serial 2-fold dilutions of peptides were added to the culture, which was then incubated for 24 hours at 37 °C at 180 rpm. The lowest antibiotic concentration that inhibited visible growth was defined as the MIC. The absorbance of the sample at 600nm was measured using a microplate reader. The MIC was also assessed for CoR and CRAB stains procured from Government Medical College, Sector 32, Chandigarh.

### 1.12 Stability of peptides

The change in MIC was assessed in presence of 150mM NaCl, 1mM MgCl_2_ and 2.5mM CaCl_2_ in order to determine the stability of peptides in a physiological environment (Ajish et al., 2022). To determine the effect of pH on the antimicrobial activity of the peptides, the pH of MH media was adjusted to 3, 5, 9 and 12 by 0.1M HCl and 0.1M NaOH (Cai et al., 2021), and the MIC was determined. The change in MIC was also determined in presence of 3% and 5% human serum, to measure degradation due to the presence of human proteases.

### 1.13 Hemolysis assay

A hemolysis assay was done to determine the toxicity of all peptides by measuring the haemoglobin release from human RBC’s. Fresh human blood was collected and washed thrice with PBS. 100ul of a 1% blood suspension was treated with different concentrations of peptide for 18h at 37°C. RBCs treated with 0.1% Triton X100 were taken as the positive control for hemolysis and untreated RBC suspension as the negative control. After completion of the incubation period, the cells were centrifuged, and the absorbance of the supernatant containing lysed erythrocytes was measured at 470 nm (Hu et al., 2016).

### 1.14 Cytotoxicity

The Cytotoxicity of AMPs against mammalian cells was evaluated using the HeLa cell line (Kashyap et al., 2022). Cells were cultured in RPMI containing 10% FBS in 5 % CO_2_ at 37°C till they reach confluence stage. Cells were detached by trypsinization and seeded into a 96-well plate containing 8-10×10^3^ cells/well in a 100ul RPMI medium. The cells were treated with AMPs at concentrations ranging from 0.5×MIC to 16×MIC and the plates were incubated for 24 h at 37°C in 5% CO_2_. Untreated cells were used as a negative control. After 24h, 0.5mg/ml of MTT reagent was added to each well and further incubated for 4 hours. The contents of the wells were removed and DMSO added to each well. Subsequently, the absorbance of each well was measured at 570nm with a micro plate reader. Cell viability was calculated using the following formula:

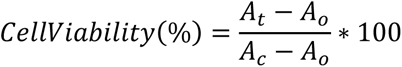

Where A_t_ is the absorbance at 570 nm of the treated cells, A_0_ is the background absorbance (at 570nm), and A_c_ is the absorbance of the negative control.

### 1.15 NPN Assay

The ability of AMPs to permeabilize the outer membrane of *A. baumannii* was determined by the quantification of fluorescence of lipophilic dye N-phenyl-1-napthylamine (NPN). Late log phase cells of *A. baumannii* 17978 were treated with 1× and 2×MIC of AMPs for 1 hour at 37°C. Cells were harvested, washed and resuspended in PBS. 10µM NPN was added to 200ul of bacterial cell suspension in a microtiter plate. Fluorescence intensity was measured immediately at excitation and emission wavelengths of 350 and 420nm, respectively. The fluorescence was normalised with the growth (O.D. _600nm_) of respective sample. Untreated *A. baumannii* cells were used as the control (Kashyap et al., 2021).

### 1.16 Membrane depolarization

The cationic, membrane-permeable dye 3, 3′-Dipropylthiadicarbocyanine iodide [DiOC_2_] was used to investigate the ability of AMPs to depolarize bacterial membranes. The membrane depolarisation activity of AMPs was measured against *A. baumannii*. The bacterial culture was washed twice in PBS containing 1% glucose and treated with 10µM DiOC_2_ for 30 minutes at 37°C in the dark. Cells were analysed using a multimode microtiter plate reader with excitation and emission intensity of 485nm and 650nm, respectively. After establishing a baseline reading, 1× and 2×MIC of each AMP was added to wells and the change in fluorescence monitored (Kashyap et al., 2021).

### 1.17 Reactive Oxygen Species (ROS) estimation

The generation of Reactive Oxygen Species (ROS) in *A. baumannii* in the presence of AMPs was estimated using 2’,7’-dichloroflourescien diacetate (DCFDA).1% overnight culture of *A. baumannii* was inoculated in 10ml LB and incubated at 37°C at 180rpm for 4 hours till the cells reach late exponential phase. Cells were washed twice with PBS and treated with 1×MIC of peptides for 1 hour at 37°C. 5µM DCFDA was added to cell suspension and incubated for 30 minutes in the dark. Fluorescence was determined with excitation and emission intensity of 488 and 530nm, respectively (Eruslanov and Kusmartsev, 2010). To evaluate the role of ROS in the antimicrobial action of AMPs, their MIC was determined in the presence of subinhibitory concentration of ROS quenchers, 200Mm thiouria and 600µM 2,2-bipyridyl (Kaur et al., 2018).

### 1.18 Checkerboard assays

The synergistic activities of peptides with different classes of antibiotics were determined using the checkerboard assay. AMPs and antibiotics were serially diluted in a 96-well microtiter plate, following the combination of two compounds in increasing concentration, and incubated in MH medium with 2×10^5^ CFU/ml of bacteria. The Fractional Inhibitory Concentration (FIC) was calculated and used to categorize the interaction of antibiotic and antimicrobial peptide. The FIC index cut-off to establish synergy was FIC ≤ 0.5. The range FIC 0.5 to 4 established an additive effect, and FIC>4 was the cut-off for antagonism (Costa et al., 2019). The FIC was calculated according to the following formula:

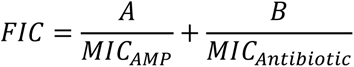

Where A is the MIC of the AMP when used in combination with the antibiotic, and B is the MIC of the antibiotic when used in combination with the AMP. MIC_AMP_ refers to the MIC of the AMP when used alone, and MIC_Antibiotic_ refers to the MIC of the antibiotic alone.

### 1.19 Field Emission Scanning Electron Microscopy (FESEM)

The ultrastructural changes in morphology of *A. baumannii* cells after treatment with combination of antibiotic and AMP (2×FIC) were examined by FESEM. The overnight grown culture was diluted 1:100 in 10 ml LB broth and grown for 4 h at 37°C, 180 rpm to obtain cells at late exponential phase. The cells were washed two times with PBS (pH 7.2) and re-suspended in PBS. This cell suspension was treated with 2×FIC of AIG-R5 and tobramycin for 4 hours at 37 °C, 180 rpm. Cells were washed twice with PBS and cell pellet was fixed with 2.5 % glutaraldehyde (prepared in PBS) at 4 °C overnight. Cells were dehydrated for 10 min in a series of graded ethanol (30-100%). The dehydrated cells were finally suspended in 70% ethanol, coated with gold particles via ion sputter and examined using FESEM (Kaur et al., 2018).

### 1.20 Biofilm inhibition assay

The biofilm inhibition activity of AIG-R5 and AIG-R5 in combination with antibiotics was determined against *A. baumannii* ATCC 17978 and CRAB isolates (P-1139 and DHA10). 5% overnight bacterial culture was incubated with AIG-R5 (1×, 2× and 4 MIC) in 96-well plate in LB-LS at 37°C for 24 hours. Biofilm cells were washed thrice with phosphate buffer saline (PBS) and fixed with methanol. The cell was stained with 0.5% crystal violet for 30min at room temperature. Biofilm was washed thrice with PBS and 33% glacial acetic acid was added to extract bound crystal violet which was quantified at 570nm using microplate reader. Biofilm inhibition was also assessed with combination of AIG-R5 and antibiotics (rifamycin, ciprofloxacin and tobramycin) (Esfahanizadeh et al., 2018).

### 1.21 Confocal laser scanning microscopy analysis

To confirm the inhibitory effect of AIG-R5 and Tobramycin on the biofilm, the fluorescent dyes SYTO 9 and propidium iodide were used. A sterile glass coverslip was used for growing biofilm of *A. baumannii* ATCC 17978 alone and in the presence of 1× and 2×FIC of AIG-R5 and tobramycin combination for 24 hours. The biofilm was stained with 5µM SYTO9 and 30µM propidium iodide (PI) for 30 min in the dark. The excess PI and SYTO9 were washed using PBS, and the cells were observed under a Confocal Laser Scanning Microscope (CLSM). The PI (red) signals were recorded at excitation and emission wavelengths of 493 and 636 nm, respectively, and the SYTO9 (green) signals recorded at excitation and emission wavelengths of 488 and 517 nm, respectively (Gautam et al., 2021).

### 1.22 Biofilm eradication assay

To determine biofilm eradication by AIG-R5, an *A. baumannii* biofilm was grown for 24h at 37°C in LB-LS medium in a 96-well microtiter plate. The pre-formed biofilm was treated with AIG-R5 at the concentrations of 1×, 2×, 4×, and 8×MIC and incubated overnight at 37°C. Planktonic bacteria were removed and biofilm cells washed three times with PBS. The attached biofilm was stained with 0.5% crystal violet as mentioned above and the absorbance was measured at 570nm using a microtiter plate reader. Biofilm eradication was also accessed with a combination of AIG-R5 and antibiotics (rifamycin, ciprofloxacin and tobramycin) (Spencer et al., 2018).

### 1.23 Data Analysis

Statistical analyses of the data were performed using GraphPad Prism 5.01 software ( raphPad Sofware Inc., CA, USA). ata was analysed using student’s t test and one-way ANOVA with Tukey’s modification. The results have been expressed as mean±standard deviation (SD) with three independent experiments. *P* value of ≤ 0.05 was considered statistically significant.

## Author Contributions

All authors jointly designed the research, evaluated the results, and wrote the manuscript.

V.T. performed wet-lab experiments. All authors have given approval to the final version of the manuscript.

## Declaration of interest

A.G. is a co-founder of AINovo Biotech Inc. and a consultant in the life science industry.

N.C. and P.S. are scientific advisors of AINovo Biotech.

## Data availability

Data will be made available upon reasonable request.

## Notes

### Competing Interest Statement

The authors have declared no competing interest.

