## Supplementary Data for "A Novel AI-Designed Antimicrobial Peptide Synergistically Potentiates Aminoglycosides against Colistin- and Carbapenem-Resistant *Acinetobacter baumannii*"

**SUPPLEMENTARY MATERIAL**

**Table S1.** MIC of peptides against *A. baumannii* in presence of sub inhibitory concentration of ROS quenchers.

| **ROS quencher** | **MIC (µM) of AIG-R1** | **MIC (µM) of AIG-R4** | **MIC (µM) of AIG-R5** |
| --- | --- | --- | --- |
| No (control) | 4 | 3 | 2 |
| 200Mm Thiourea | 4 | 3 | 2 |
| 600µM 2,2’-Bipyridyl | 4 | 3 | 2 |
| 200Mm Thiourea + 600µM 2,2’-bipyridyl | 4 | 3 | 2 |

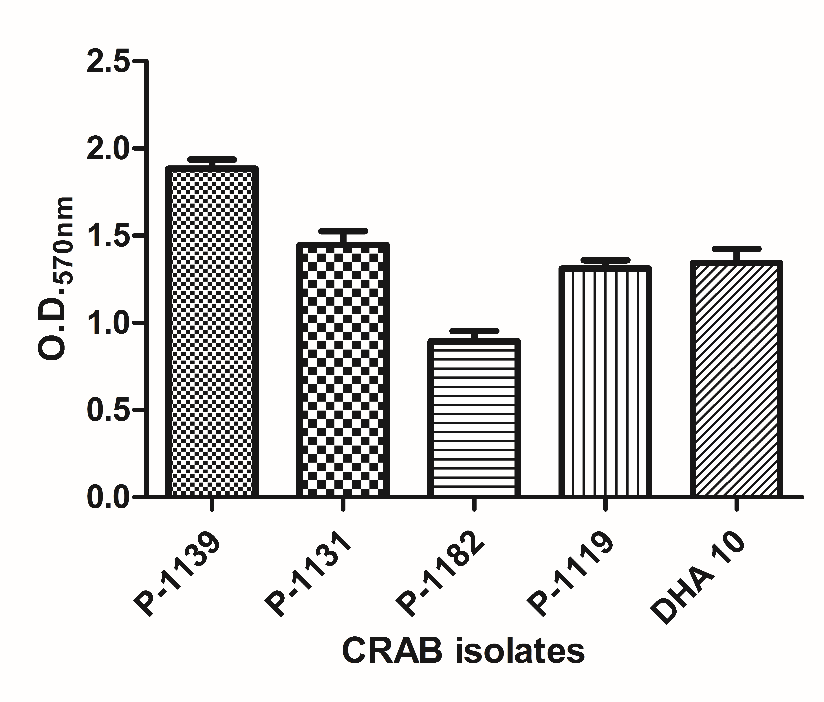

**Figure S1.** Biofilm formation by CRAB isolates. The results are expressed as mean ± SD of three independent experiments.

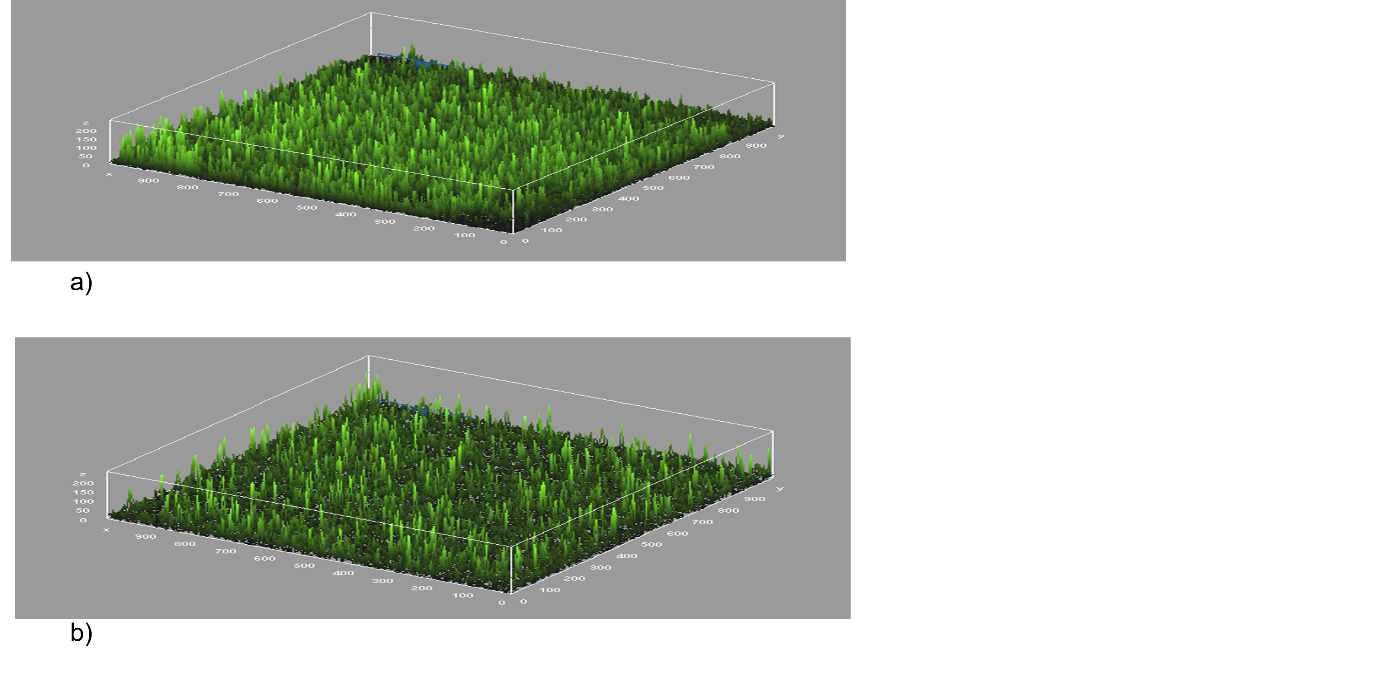

**Figure S2**. Three dimensional surface representation of *A. baumannii* biofilm (a) Untreated and (b) 1/8×AIG-R5 and 1/4×tobramycin treated biofilm.

**Table S2**. Antibiotic resistance profile of *A. baumannii* isolates

| A. ***baumannii* Strains** | **Imipenem** | **Meropenem** | **Colistin** |
| --- | --- | --- | --- |
| **17978** | S | S | S |
| **Ci 3** | R | R | R |
| **Ci 6** | R | R | R |
| **Ci 7** | R | R | R |
| **P-1139** | R | R | R |
| **P-1182** | R | R | R |
| **P-1119** | R | R | R |
| **P-1131** | R | R | R |
| **P-1169** | R | R | R |
| **DHA 8** | R | R | R |
| **DHA 10** | R | R | R |

S= sensitive; R= resistant

**Table S3**. Stability of AMPs at different pH levels.

| **AMPs** | **MIC (µM) against *A. baumannii* ATCC 17978** | | | | |
| --- | --- | --- | --- | --- | --- |
|  | **pH 3** | **pH 2** | **pH 7** | **pH 9** | **pH 12** |
| **AIG-R1** | 4 | 4 | 4 | 9 | 12 |
| **AIG-R4** | 3 | 3 | 3 | 6 | 9 |
| **AIG-R5** | 2 | 2 | 2 | 2 | 2 |

**Cytotoxicity and hemolytic activity of AIG-R5 and tobramycin combination**

The combination of AIG-R5 and tobramycin at 1×FIC (1/8 AIG-R5 and 1/4 tobramycin) and 2×FIC (1/4 AIG-R5 and 1/2 tobramycin) was not cytotoxic to HeLa cells as the cell survival was 100% and 97%, respectively (Figure 4) and showed negligible hemolysis (<0.05%) at 1×FIC and 2×FIC (Supplementary Table S3).

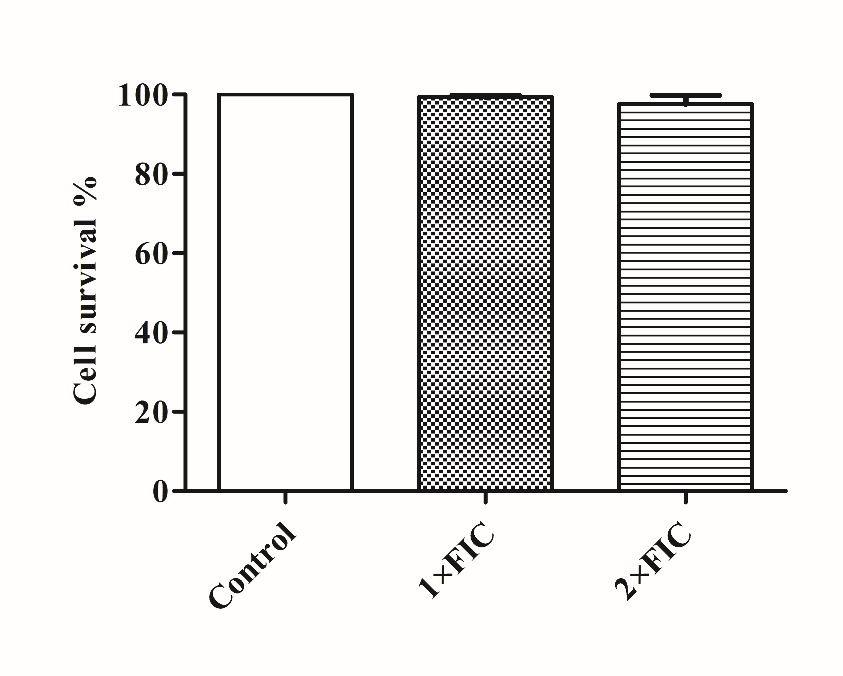

**Figure S3**. Cytotoxicity of combination of AIG-R5 and tobramycin at 1×FIC and 2×FIC against HeLa cell line. All values expressed as the mean ± SD.

**Table S4.** Hemolytic activity of AIG-R5 with tobramycin at 1×FIC and 2×FIC against human RBCs.

| **AIG-R5 and tobramycin combination** | **% Hemolysis** |
| --- | --- |
| 1×FIC (1/8×AIG-R5 + 1/4×tobramycin) | 0.018 |
| 2×FIC (1/4×AIG-R5 + 1/2×tobramycin) | 0.005 |

Values expressed as an average of three independent experiments.
